# Annotating Metagenomically Assembled Bacteriophage from a Unique Ecological System using Protein Structure Prediction and Structure Homology Search

**DOI:** 10.1101/2023.04.19.537516

**Authors:** Henry Say, Ben Joris, Daniel Giguere, Gregory B. Gloor

**Affiliations:** University of Western Ontario, Schulich School of Medicine and Dentistry, Department of Biochemistry, London ON, Canada

**Author notes:** Address correspondence to Greg Gloor,. H.S collected the samples, performed the experiments and analysis, and contributed to the design of the study. B.J. conceived and developed the secondary assembly pipeline used in the analysis, and provided input to initial assembly and polishing methods. D.G. collected and sequenced samples GAC0 and GAC1, as well as provided input and methodology development to the initial assembly and polishing methods. G.G. supervised the project and conceived the study. H.S and G.G wrote the manuscript, with revisions from B.J and D.G. Second Author, Full Affiliation.

**Keywords:** Annotation, protein structure prediction, granulated activated carbon, naphthenic acids, metagenome

## Abstract

Emergent long read sequencing technologies such as Oxford’s Nanopore platform are invaluable in constructing high quality and complete genomes from a metagenome, and are needed investigate unique ecosystems on a genetic level. However, generating informative functional annotations from sequences which are highly divergent to existing nucleotide and protein sequence databases is a major challenge. In this study, we present wet and dry lab techniques which allowed us to generate 5432 high quality sub-genomic sized metagenomic circular contigs from 10 samples of microbial communities. This unique ecological system exists in an environment enriched with naphthenic acid (NA), which is a major toxic byproduct in crude oil refining and the major carbon source to this community. Annotation by sequence homology alone was insufficient to characterize the community, so as proof of principle we took a subset of 227 putative bacteriophage and greatly improved our existing annotations by predicting the structures of hypothetical proteins with ColabFold and using structural homology searching with Foldseek. The proportion of proteins for each bacteriophage that were highly similar to known proteins increased from approximately 10% to about 50%, while the number of annotations with KEGG or GO terms increased from essentially 0% to 15%. Therefore, protein structure prediction and homology searches can produce more informative annotations for microbes in unique ecological systems. The characterization of novel microbial ecosystems involved in the bioremediation of crude oil-process-affected wastewater can be greatly improved and this method opens the door to the discovery of novel NA degrading pathways.

**IMPORTANCE:** Functional annotation of metagenomic assembled sequences from novel or unique microbial communities is challenging when the sequences are highly dissimilar to organisms or proteins in the known databases. This is a major obstacle for researchers attempting to characterize the functional capabilities of unique ecosystems. In this study, we demonstrate that including protein structure prediction and homology search based methods vastly improves the annotation of predicted genes identified in novel putative bacteriophage in a bacterial community that degrades naphthenic acids the major toxic component of oil refinery wastewater. This method can be extended to similar genomics studies of unique, uncharacterized ecosystems, to improve their annotations.

**P**lease read the [Instructions to Authors](https://journals.asm.org/journal/msystems/submission-review-process) carefully, or browse the [FAQs](https://journals.asm.org/journal/msystems/faq) for further details.

## INTRODUCTION

Next-generation long-read sequencing technologies have made it easier to generate complete metagenomically assembled genomes (MAGs) of difficult to assemble species, which is an essential first step for comprehensive investigations into whole microbial communities (1). However, even with long-read technologies it can be difficult to assemble high quality whole genomes from metagenomic samples, and these problems are even more acute when characterizing metagenomes from environmental sites. There are a number of issues that needed to be addressed in parallel to be able to extract useful information when characterizing these metagenomes. First, it is crucial to extract pure and intact high molecular weight DNA. Second, sufficient data must be collected from the sequencing run to be able to characterize species at non-trivial relative abundances. Finally, care must be taken before and after the assembly to ensure that redundant sequences and low quality reads are excluded from the assembled genomes. These considerations must be well accounted for to produce high quality sequences as they the foundation of subsequent analyses - including genome annotation.

Although many tools and techniques exist to predict the organization and functional capabilities of MAGs, they fundamentally base their annotations directly on DNA or proteins sequences. This is an issue when sequences of interest appear to be completely novel, and it is common for a majority of predicted genes in MAGs to end up unannotated since they are below the threshold for homology finding by sequence alignment (2). This is especially the case for the samples sequenced in this study, in which we use long-read sequencing to assemble and annotate genomes to characterize a unique microbial ecosystem that grows in naphthenic acids (NAs) enriched environment. NAs are the major toxic contaminant of oil refinery wastewater.

There would be many benefits to functionally annotating ecological systems such as this. In the past, other bacterial species living in oil processing and refining wastewater that degrade a small number of surrogate NAs have been observed, but no single organism or mixture of organisms has been identified, sequenced or annotated that can efficiently degrade the full spectrum of NAs (3). One reason for this may be because the isolates tend to be identified from environments that are highly heterogeneous, such as tailings ponds, where NAs are a minor constituent and generally a carbon source of last resort (4). Furthermore, the heterogeneity of environmental samples make it difficult to isolate naturally occurring NAs, necessitating the use of surrogate NAs when enriching environmental samples for NA degrading bacteria to study (3). However, this study’s samples of interest originate from an oil refinery waste-water treatment plant where the last step of its multi-step treatment process involves the capture of residual NAs on industrial-scale granulated activated carbon (GAC) beds. This provides a unique environment highly enriched for NAs, upon which a bacterial biofilm grows using the NAs a carbon source. The biofilm initially aids in the remediation of wastewater from NAs, but ultimately overgrowth fouls the GAC beds necessitating frequent and regular exchange. As such, the samples collected in this study presents a unique opportunity to investigate and annotate NA degrading bacteria as it serves as a natural experiment. Functionally annotating this particular bacterial community would be highly informative for any future development of bioengineered ecosystems for ecological treatment and restoration of oil contaminated sites, which is in the best interest of the industry and environment (5).

Given the complexity and novelty of this microbial community, it is unsurprising that the use of state of the art sequence based homology annotation pipeline, such as Bakta (6), provided poor annotations because of the sequence novelty. However, since protein structures are often more conserved than is sequence and that protein structures are often highly correlated to their functions, the inclusion of structure based homology into the analysis was expected to substantially enrich our annotation ability by detecting remote functional relationships (7). Advances over recent year in the efficiency, accessibility, and accuracy for computational methods of predicting protein structures and performing structural homology searches, such as with Colabfold (8) and Foldseek (9) made querying a large number of putative coding sequences for proteins from many genomes in a reasonable time frame feasible.

Here we present wet- and computational-lab approaches that allow the sequencing, assembly and annotation of high quality assembled circular contigs (ACCs) from GAC samples. We identify ACCs of sub-genomic size and characterize their bulk properties. Furthermore, we extract a subset of sub-genomic ACCs that are putative bacteriophage. Given current limitations of computational resources needed for protein structure prediction of an entire metagenome, our current focus for structure based annotations shifted to our subset of putative bacteriophage to demonstrate proof of principle. We show that structure-based annotation based on ColabFold (8) and FoldSeek (9) was able to add substantial annotation power, even for bacteriophage which are notoriously hard to annotate.

## RESULTS AND DISCUSSION

Samples of the microbial biofilm growing on the granulated activated carbon (GAC) beds were collected from the refinery’s wastewater treatment plant between 2019 and 2022. Table 1 summarizes the collection dates, sequencing and assembly information. Initial metagenomic characterization of the microbial ecosystems with shotgun Illumina sequencing showed that the species found were essentially unique, and that an alternative approach to characterization and annotation would be required. For this reason, we took advantage of the long-read Oxford Nanopore technology and its rapidly improving DNA sequence read quality.

**TABLE 1.** Sequencing Statistics of the 10 GAC samples collected from the oil refinery wastewater treatment facility. All samples were sequenced using the Oxford Nanopore MinION platform. The read statistics represent the pre-filtered metagenomic reads used for the assembly pipeline, and the assembled circular contigs (ACCs) listed represent primary + secondary assemblies that are polished and circularized.

| Sample | Date Collected | Gbs | Mean Q-Score | rN50 (kbp) | ACCs (P+S) | Putative bacteriophage |
| --- | --- | --- | --- | --- | --- | --- |
| GAC0 | xx/xx/2019 | 14.54 | 13.5 | 8.1 | 195 + 101 | 6 |
| GAC1 | 27/01/2020 | 9.88 | 13.8 | 4.5 | 314 + 31 | 10 |
| GAC2 | 14/02/2020 | 19.96 | 13.6 | 20.0 | 453 + 76 | 28 |
| GAC3 | 02/03/2020 | 22.71 | 14.3 | 17.9 | 624 + 114 | 38 |
| GAC4 | 06/10/2020 | 26.65 | 13.2 | 12.4 | 405 + 69 | 9 |
| GAC5 | 03/05/2021 | 15.06 | 13.3 | 12.3 | 561 + 98 | 38 |
| GAC6 | 07/05/2021 | 19.95 | 13.4 | 9.5 | 405 + 72 | 18 |
| GAC7 | 14/05/2021 | 24.40 | 13.3 | 6.4 | 746 + 110 | 37 |
| GAC8 | 10/08/2021 | 35.20 | 14.5 | 30.1 | 445 + 38 | 18 |
| GAC9 | 21/04/2022 | 33.52 | 13.6 | 9.7 | 494 + 81 | 25 |

Gbp: giga-bases of raw sequence, Mean Q-score: mean quality score of raw sequenced reads, rN50: raw read N50

The first sample, GAC0, was collected in 2019 but the day and month was not noted. Ultra-high molecular weight DNA was isolated from these samples by gentle cell lysis optimized for the high heavy metal content in the environment. This was followed by spooling from an aqueous/ethanol interface, subsequent clean-up, and size selection for higher molecular weight fragments as described in the Methods section. Purified DNA was sequenced on the Oxford Nanopore Technologies (ONT) MinION platform, base-called and assembled as outlined in the Methods and summarized in Figure 1. Metagenomically assembled fragments less than 1 Mb that had properties consistent with fully closed, non-repetitive circular DNA sequences with an estimated coverage of greater than 10 were subset and polished; note that circularly permuted sequences would also appear to assemble as circular DNA fragments. In the end, we had 5432 polished ACCs across the 10 samples for analysis and a total of 227 putative bacteriophage.

**FIG 1.**
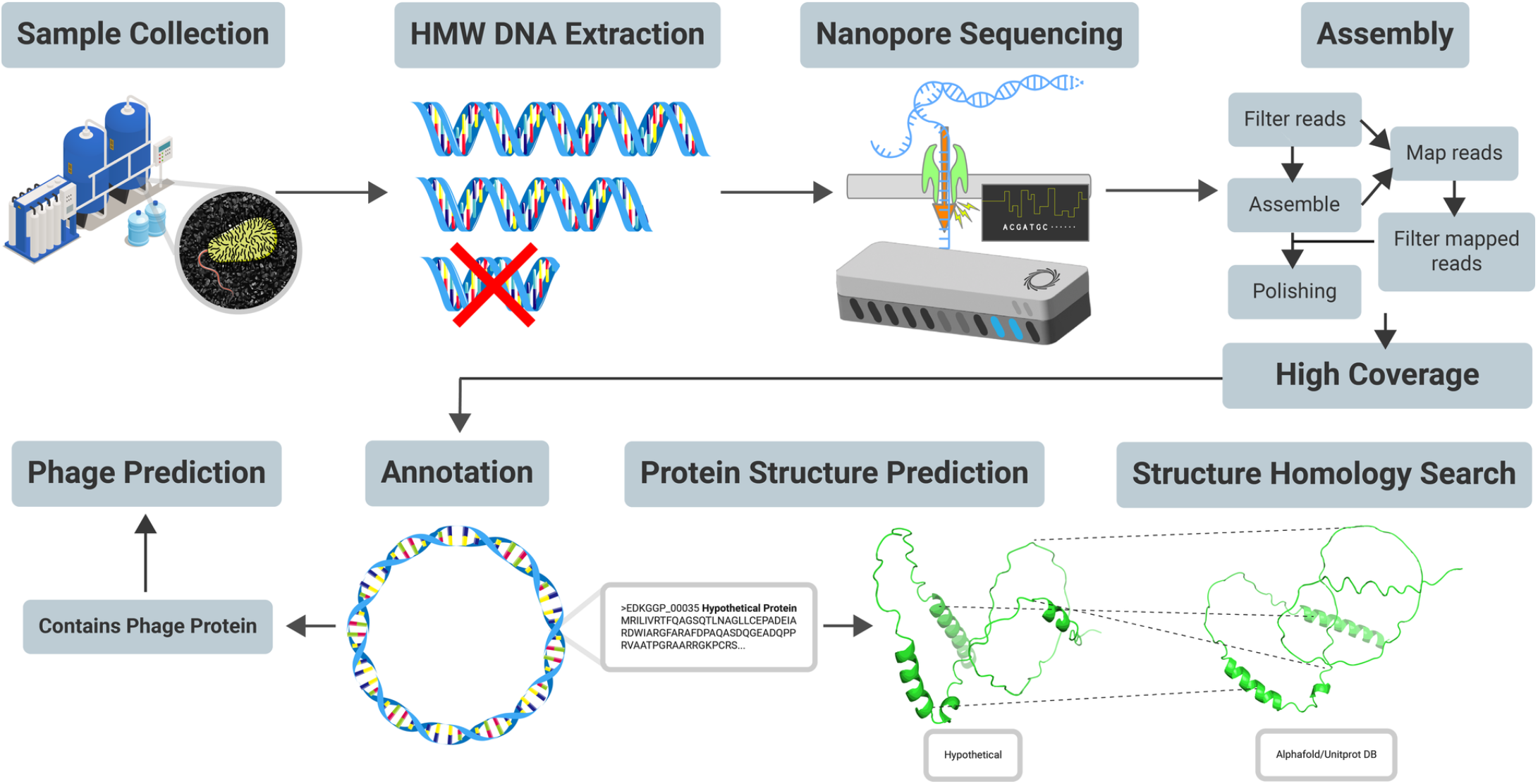
Samples were collected from the GAC beds at an oil refinery site in Ontario when the beds were being exchanged. Ultra-high molecular weight DNA was extracted and low molecular weight fragments were removed by differential PVP precipitation. A library was made from the purified DNA using the Oxford Nanopore LSK109 or LSK110 kit, and was sequenced on the Oxford Nanopore MinION platform; an entire 9.4.1 flow cell was used for each sample. The DNA was assembled using a custom workflow that included a post assembly filtering step using gerenuq (10) to remove low-quality and poorly mapped reads from the assembly. Only fully closed circles with an estimated minimum coverage of 10 were kept for further analysis. Initial annotation was done using Bakta (6) and ACCs that had any open reading frame annotated as a bacteriophage protein were kept for both prediction and structural annotation via ColabFold (8) and FoldSeek (9).

We examined the size distribution of the ACCs less than 1 Mbp and Figure 2A shows a density plot of all sub-genomic ACCs across all samples. This shows only those < 100 kbp since there was essentially no density in the interval between 100 and 1000 kbps. We observed that each sample produced assembled fragments that shared a bimodal size distribution with the highest densities at approximately 10 and 42 kbps, with a smaller peak at 60 kbps. We next examined the size distribution of ACCs seen once, and those seen multiple times. For this, an all versus all BLAST comparison was performed to identify any ACCs that occur in multiple samples by finding reciprocal best hits with a percent identity of 98%, a query coverage that was within 10% of the query length, and a query length that was at least 90% of subject length. We observed that the ACCs that appear in either only one sample, which represent the majority, and recurring ACCs shown in Figure 2B and C, share a similar size distribution.

**FIG 2.**
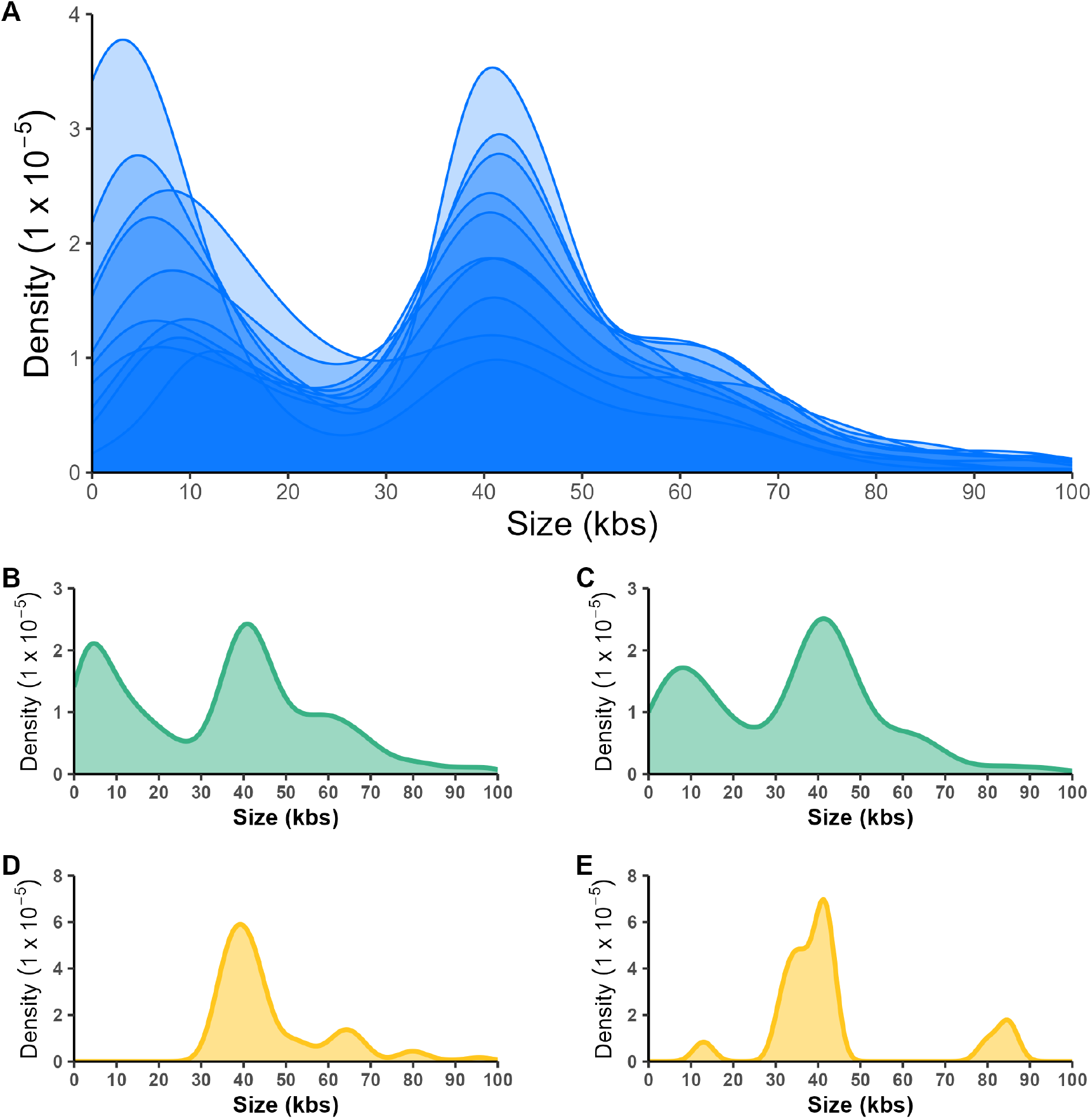
Size distributions of assembled circularized contigs (ACCs) obtained from the metagenomes of 10 GAC samples (A) Each density curve represents the subset of draft assemblies under 100 kbp that circularized in each of the 10 GAC samples. Each curve follows a bimodal distribution across the 10 GAC samples that were sequenced, with a majority of contigs in this range appearing at approximately 8 and 42 kbp. (B) ACCs that appeared only in 1 sample follows a similar bimodal distribution. The majority of polished assemblies fall under this category. (C) A subset of polished ACCs that appeared in more than 1 sample follows the same bimodal distribution but with a hihger density at the 42 kbp size. (D) The distribution of predicted bacteriophage ACCs that only appeared in a single sample differed in that bacteriophage in this subset were mainly 40 kbp and up. (E) Predicted bacteriophage that appeared in more than one sample were mainly those in the 42 kbp range, but also in the 85 and 12 kbp ranges.

Polished ACCs were annotated using Bakta (6) and 423 ACCs containing at least one putative bacteriophage related protein were passed on to the INHERIT package (11) which assigns scores based on the inferred likelihood of being bacteriophage; of these, 227 bacteriophage were predicted. The subset of 227 circularized ACCs that contained at least one putative bacteriophage protein annotation by Bakta and that were predicted to be bacteriophage by INHERIT demonstrated different size distributions as shown in Figure 2D and E for those observed in one sample and those observed in multiple samples. The greatest density of putative bacteriophage in the single sample category was approximately at 40 kbp, with smaller peaks at 65, 80 and 95 kbp. The size distribution of putative bacteriophage in the recurring category had a major density peak at 42 kbp, with smaller peaks at 12, 35 and 85 kbp.

The consistency in size distribution for most subgenomic sized ACCs across samples could either be due to a particular set of selectively advantageous genes, or even type of plasmid mobility. Previous studies that have also observed bi-modal distribution of plasmids within species observed correlations between mobility type and plasmid sizes (12). Additionally, it is possible that size constraints can result from bacteriophage transduction of plasmids which are limited by size compatibility and similar to the observation of bacteriophage and plasmid size distributions made in the study by (12). Interestingly, the largest peak in our putative bacteriophage ACCs matches the largest of the two peaks observed for all ACCs at approximately 42 kbp. Though it is not yet clear, this may be a mechanism that explains the bimodal distribution of ACCs up to 100 kbps seen here.

Standard metagenomic quality control metrics are not available for the predicted bacteriophage making it impossible to assess the quality of the circularized assemblies. Instead, we assessed assembly quality by examining the relationship between coverage depth and uniformity to see if the putative bacteriophage ACCs were similar in coverage and uniformity of coverage to high quality MAGs. Assessing the quality of MAGs follows a well-defined set of ruled regarding genome coverage, gene content and gene redundancy (13). For this, we collected circular MAGs of > 1 Mbp with > 90% completeness and < 10% contamination from this dataset, which were assembled and polished using the same methods used for the bacteriophage ACCs. We collected 37 high quality MAGs with mean coverage between 5 and 1000 and plotted the per-position mean coverage and the per-position standard deviation as shown in Figure 3A. We observed a linear relationship as expected, with the SD/cov ratio being less than 1 for all high quality MAGs. We did the same calculations for the ACCs and found that the vast majority also had an SD/cov ratio less than one showing that the majority of these were consistent with the coverage characteristics observed for high quality MAGs. However, there was a subset of bacteriophage assemblies that have higher than expected SD/cov ratios.

**FIG 3.**
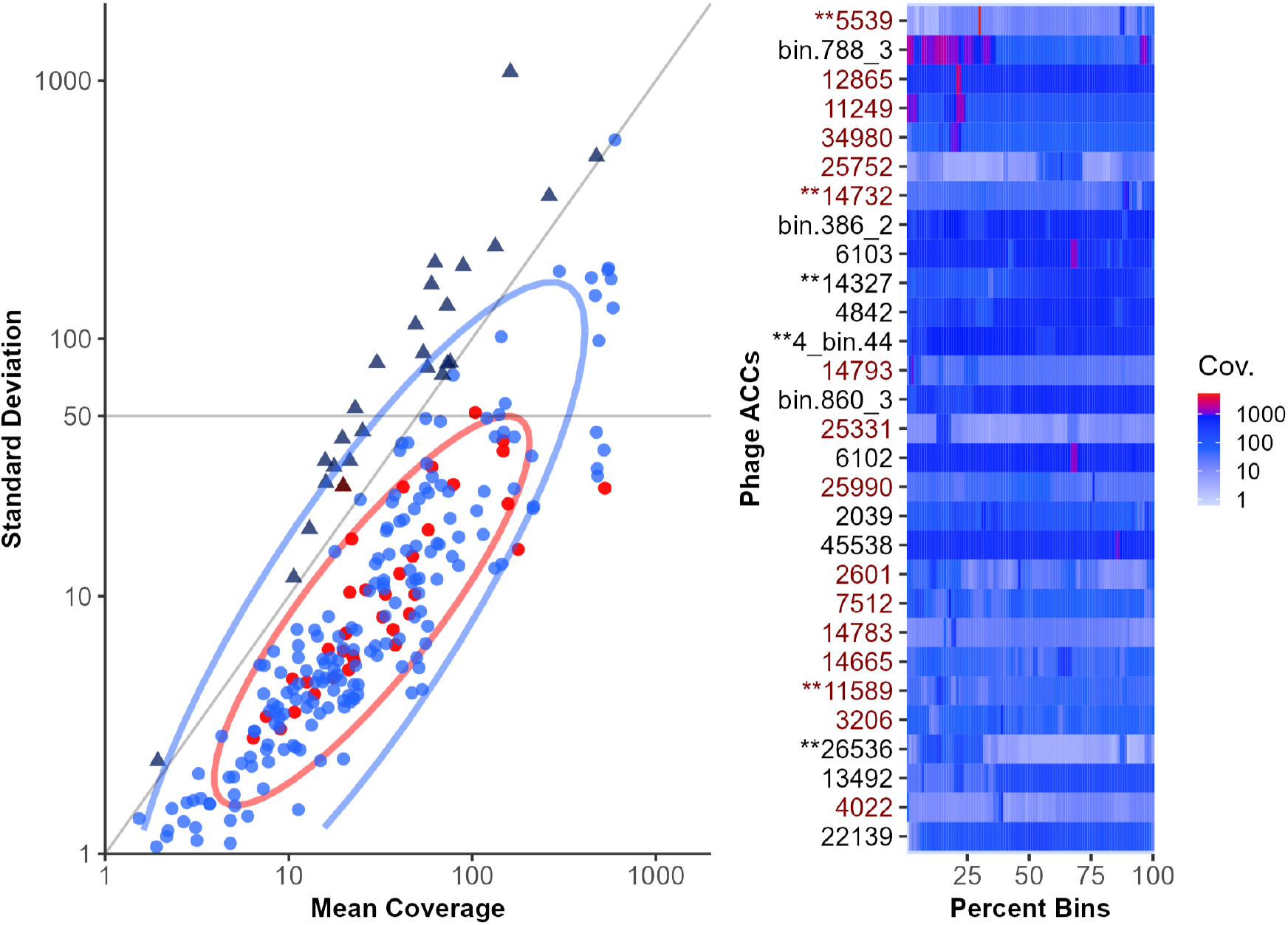
Evaluating the quality of ACC assembly using a coverage-based metrica metric. Panel A shows the standard deviation (SD) of the per-base coverage (cov) plotted as a function of the mean per-base coverage. This is shown for high quality MAGs (red) which were at least 90% complete and less than 10% contaminated and for predicted bacteriophage ACCs (blue). Both were assembled from the GAC samples with the same pipeline detailed in the Methods section. The ovals show the 90% confidence interval of each group and reveal that although a majority of predicted bacteriophage ACCs followed a trend consistent with high quality MAGs, a number of them had standard deviations higher than expected. Bacteriophage ACCs with a SD/cov ratio greater than 1 are represented by the triangle symbol. Panel B shows a plot of the mean coverages over 1% bins for those putative bacteriophage ACCs with standard deviations higher than 50; these are ordered from highest SD to lowest. This heatmap visualizaton helps identify potential problematic regions that lead to high standard deviations and inflated mean coverages. Most bacteriophage ACCs in this subset have one or more coverage spikes that span 1-5% of the genome. Conversely, there are few bacteriophage ACCs with relatively even coverage for most of the genome but have drops in coverage over broad regions. ACCs denoted with ** contain at least one read that aligns over the full length with at least 90% sequence identity to the reference. Labels in red indicate those ACCs with SD/cov ratios greater than 1.

Figure 3B shows a closer examination of bacteriophage ACCs with standard deviations over 50. Here the ACCs are ordered from highest to lowest ratio, and shows that many ACCs in this subset have small regions that have much higher mean coverages leading to a high standard deviation of the coverage for the ACC. These could represent hybrid assemblies between closely related bacteriophage, direct repeats, or circular permuted sequences where the end sequences were not perfect direct repeats. In addition to regions of abnormally high coverage, some assemblies have drops in coverage over broader regions. Lower coverage regions may be a result of the depletion of smaller DNA fragments during the DNA extraction and sequencing library prep steps, which was originally done since bacterial genomes and plasmids were the initial goal. Regardless, for some assemblies highlighted in Figure 3B such as ACC 5539, there exists at least 1 full length read alignment with a minimum 90% sequence identity that can be used to provide some validity to the assembly. However, for other assemblies it is not yet clear if the coverage spikes or drops indicate misassemblies or not. We concluded the majority of the ACCs of putative bacteriophage were assembled and polished to a standard consistent with that observed for high quality MAGs. Furthermore, in the absence of other information, the SD:coverage ratio can be used as a proxy to help identify poorly assembled sub-genomic circles.

Characterizing these predicted bacteriophage on a functional level was difficult as a majority of the predicted coding sequences (CDS) annotated by Bakta (6) remained hypothetical. To create more informative annotations and identify many of these hypothetical CDS, predicted CDS from Bakta were passed to Colabfold (8) to predict protein structures and then to assess structural homology vs. the entire universe of known and predicted protein structures using Foldseek (9). Structural homology searching is much more sensitive than sequence homology searching as only the fold and not the sequence needs to be conserved (7).

Using this approach, a substantial increase in identifiable proteins and annotations with GO or KEGG terms was observed for all CDS in all predicted bacteriophage ACCs across all samples when compared to sequence homology annotation alone (Figure 4). We observed that Bakta annotations using PFAM(14) provided an annotation for about 10-15% of CDS in all samples, while the structural homology search approach gave confident predictions for between 50 and 75% of CDS. We found that annotation of CDS with KEGG (15) and GO terms (16) was particularly under-represented in Bakta annotations, with the majority of CDS having no annotated function. In contrast, about 25% of CDS were able to be annotated with a KEGG or GO functional term. Indeed, the combination of Colafold and Foldseek substantially outperformed sequence only annotation by Bakta in this aspect, producing at least 1 or more KEGG or GO terms for every predicted bacteriophage across all samples.

**FIG 4.**
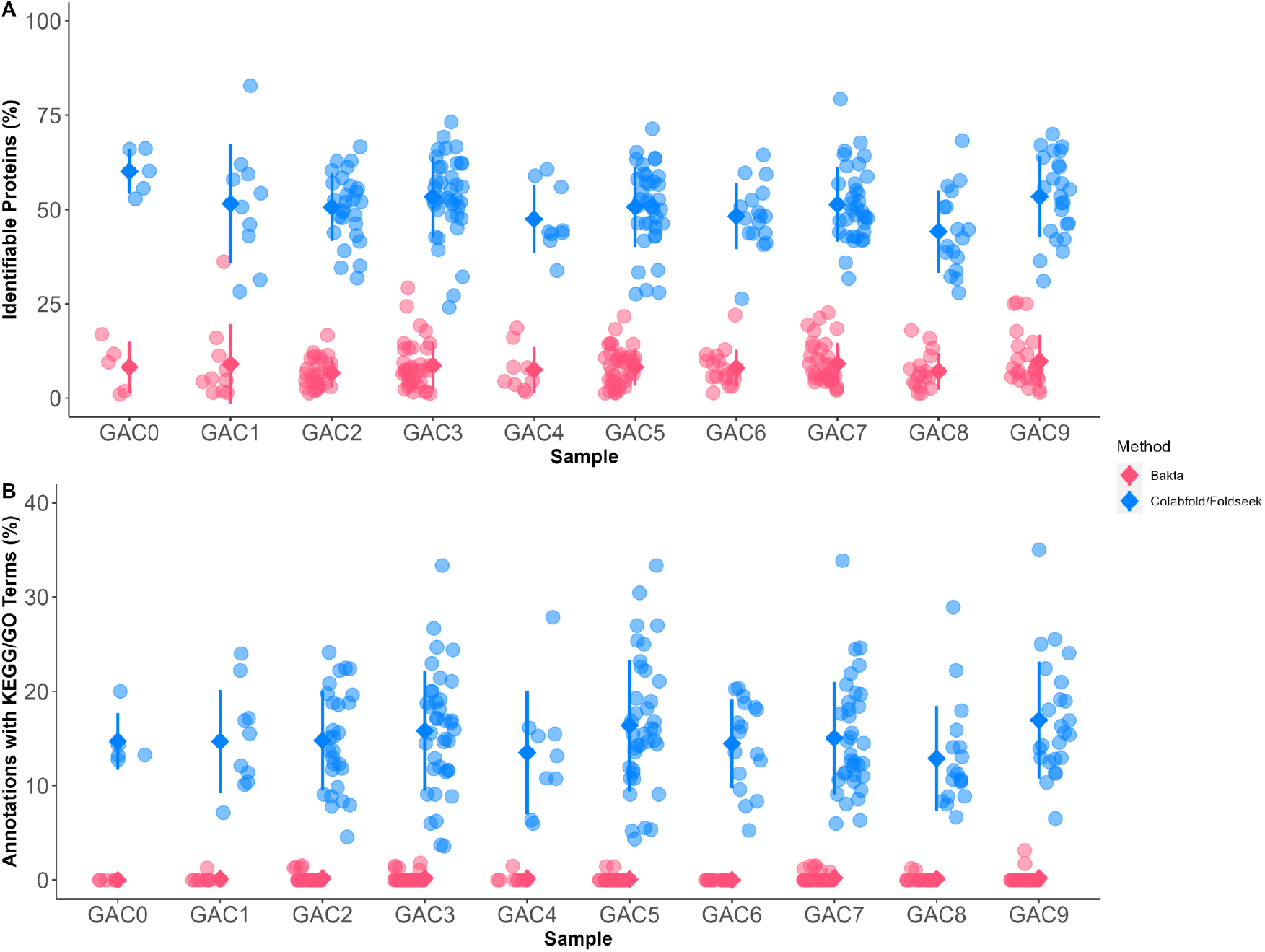
Using protein structure prediction and structure homology search methods (Colabfold and Foldseek) consistently increases the proportion of predicted CDS’ that are annotated in bacteriophage across all samples versus sequence homology alone (Bakta). (A) A large proportion of hypothetical proteins identified by PFAM annotations were able to be identified using Colabfold and Foldseek. (B) The proportion of annotations that also include KEGG or GO terms increased consistently across all predicted bacteriophage across each sample. Whereas Bakta annotations had no KEGG/GO terms for most bacteriophage ACCs, Foldseek led to KEGG or GO annotations in all bacteriophage ACCs.

This result allowed us to identify a number of functions and pathways present in putative bacteriophage ACCs in this sample, many of which were functions and pathways expected to be found in bacteriophage Figure 5. The most common GO terms included processes related to DNA metabolism, repair, recombination and replication processes, in addition to host infection or interaction such as hydrolase activity or membrane related activity. KEGG pathways that appeared in bacteriophage ACC Foldseek annotations also included expected pathways in most samples, including homologous recombination and mismatch repair pathways. Metabolically, the number of GO terms produced were diverse but their relation to naphthenic acid degradation was not clear. The carboxylic acid and lipid metabolic processes however, may be naphthenic acid related as previous proposed mechanisms of naphthenic acid degradation involved *α* -oxidation or *β* -oxidation which are reactions included in carboxylic acid or lipid metabolism (4). Regardless, many of these observations regarding the functional capabilities of our bacteriophage ACCs would not have been possible with a sequence homology only approach.

**FIG 5.**
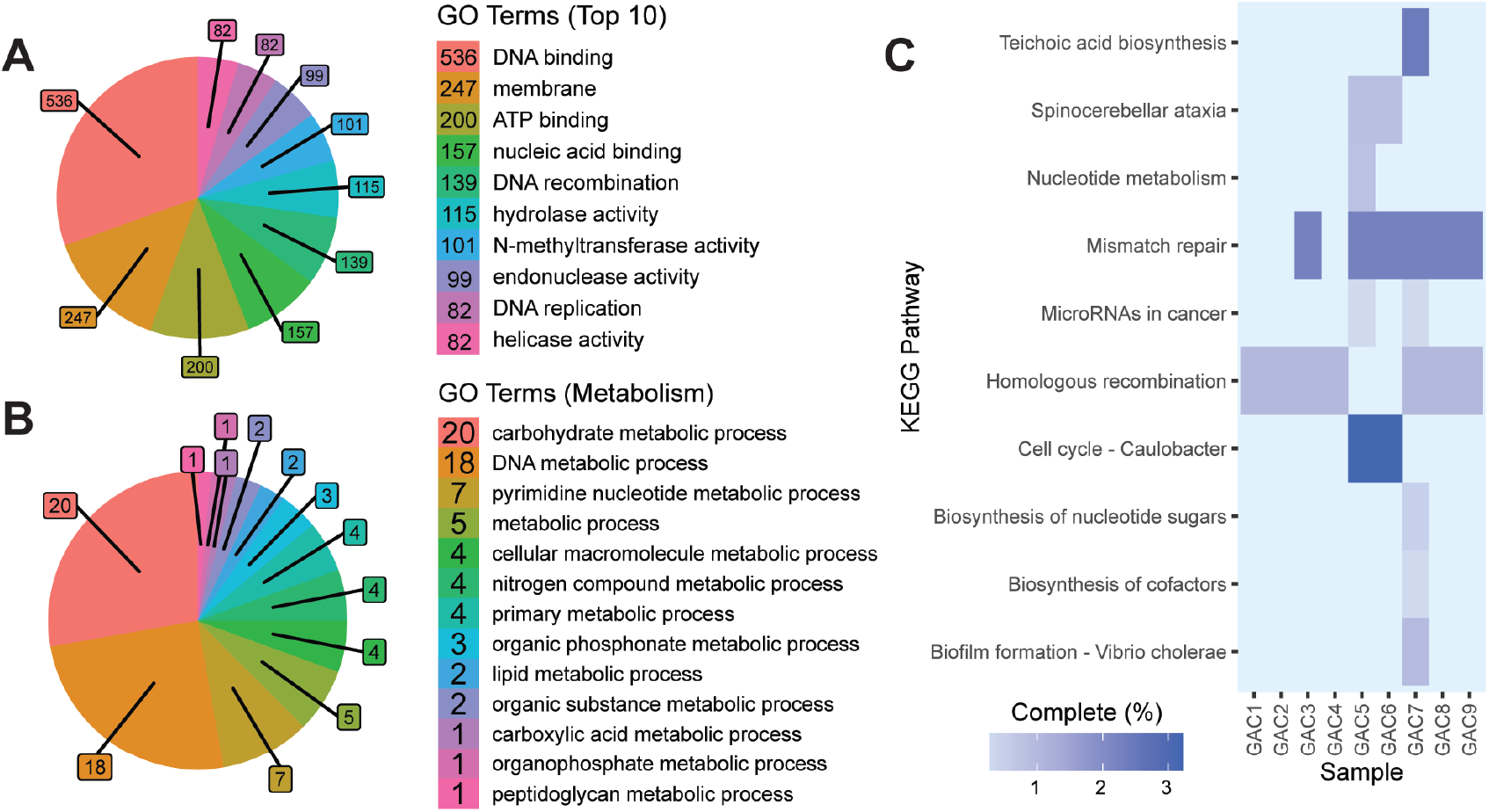
Foldseek revealed many functional pathways and processes present in putative bacteriophage ACCs, many of which were expected bacteriophage pathways. (A) The top 10 GO terms across all putative bacteriophage included mainly GO terms relevant to the bacteriophage life cycle, including DNA related functions and functions related to host infection and interaction. (B) Proteins involved in a variety of metabolic processes were identified across all the putative bacteriophage. Though carboxylic acid and lipid metabolic processes represent the minority, they may potentially be related to NA degradation based on past proposed mechanisms of NA degradation. (C) The presence of KEGG pathways were identified from Foldseek annotations for the entire set of predicted protein structures in all putative bacteriophage in each sample. Some pathways common to bacteriophage, including mismatch repair and homologous recombination, were expected and present in most samples. All the pathways annotated in our samples were incomplete KO’s in putative bacteriophage Foldseek annotations across each sample represented only a small fraction of total unique KO’s from each KEGG pathway.

The structural based annotation approach identified more putative homologs than sequence based annotation for all size classes of CDS, with the greatest number of putative homologs being identified in CDS between 100 and 500 bp. The sequence homology approach was able to annotate up to approximately 25% of predicted CDS’ up to 5 kbp in size, whereas the structural homology method was able to annotate from approximately 25% to 80% of CDS’ up to 6 kbp in size. Examining this data as proportions reveals that the structural homology approach was particularly efficient for CDS greater than 500 bp, where between 60 and 75% of CDS between 0.5 and 4 kbp were annotated by this approach. CDS identification using Bakta predicted CDS of up to 12 kbp in size, although any CDS greater than 5 kbp in size remained hypothetical, whereas over 25% of CDS in this size range were able to be annotated structurally (Figure 6). However, a the vast majority of predicted CDS that were below 0.5 kbp remained unannotated by either method. Thus, the small proportion of identifiable proteins relative to the number of proteins identified in that size bin reduce the overall frequency to about 50% across all size ranges.

**FIG 6.**
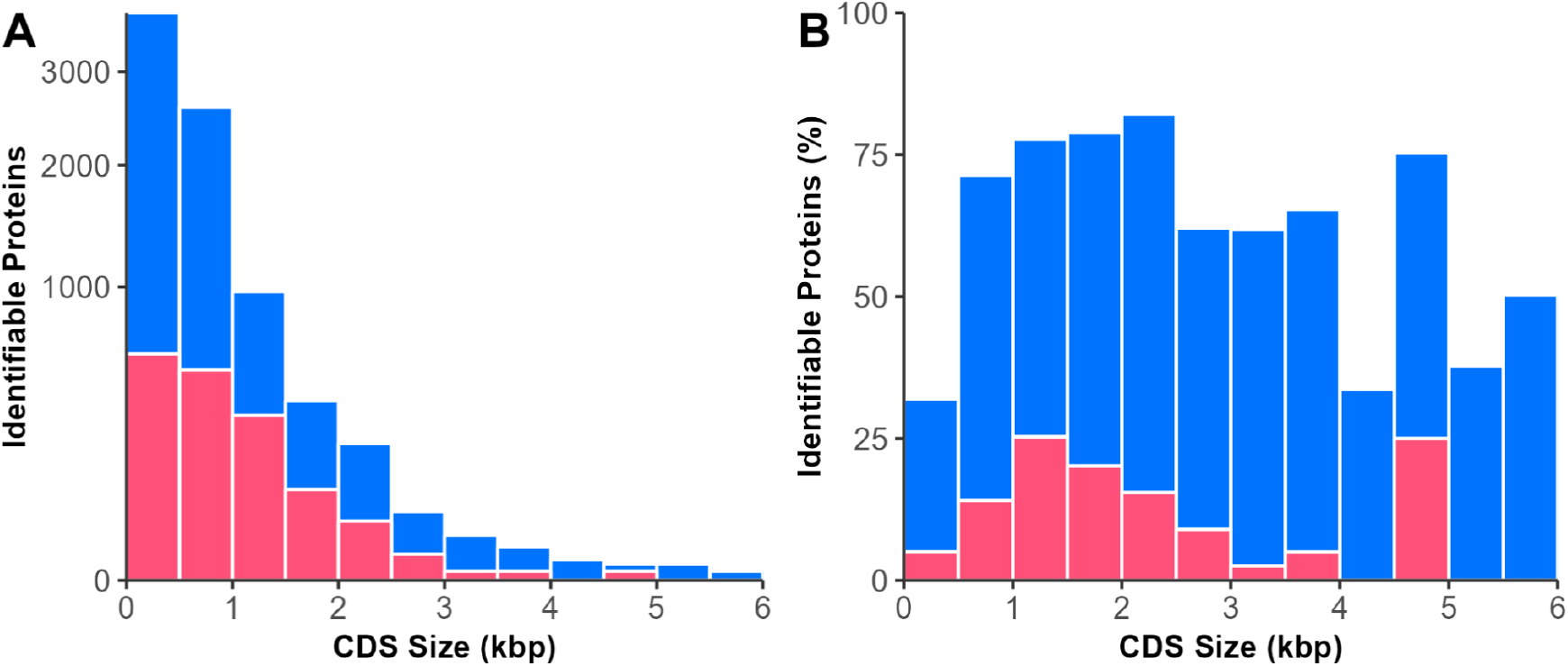
Including protein structure predictions and homology search based methods result in a larger proportion predicted CDS’ of up to 6 kbp being annotated in predicted bacteriophage. (A) There is a universal increase in the proportions of CDS’ annotated of sizes up to 6 kbp when using structure homology. CDS’ between 5 kbp to 6 kbp that were not annotated with sequence homology methods were able to be annotated using structure homology. CDS’ between 6 and 12 kbp were also predicted by Bakta but could not be annotated with either method. (B) The number of CDS’ predicted and annotated is related to CDS size. While the number of identifiable proteins more than doubled when including structure homology methods for CDS’ of sizes less than 5 kbp, an overwhelming proportion of proteins from CDS less than 0.5 kbp CDS size range remained unannotated.

We examined the concordance between PFAM annotations output by the sequence and structural homology search approaches including data from all samples, and show the results in Figure 7A. We found that across all CDS with an annotation by either the sequence or structural homology approach, ColabFold and Foldseek together was able to identify 583 PFAM annotations whereas Bakta found only 296, with 183 (26%) of these being in common. Thus, the total PFAM identifications found by the structural approach was not a strict superset of all of PFAM identifications found by sequence homology. A more rigorous way of comparing the annotation methods was to examine the overlap when both approaches provided an annotation for the same CDS shown in Figure 7B. Of the 194 CDS where both approaches predicted a CDS, we observed that 141, or 72%, were concordant for PFAM identifications. In the absence of orthogonal evidence, we were unable to determine which approach was more accurate, but given the extremely high sequence divergence between the data collected here and the sequence databases it is possible that many of the sequence-only annotations may be spurious or of low confidence.

**FIG 7.**
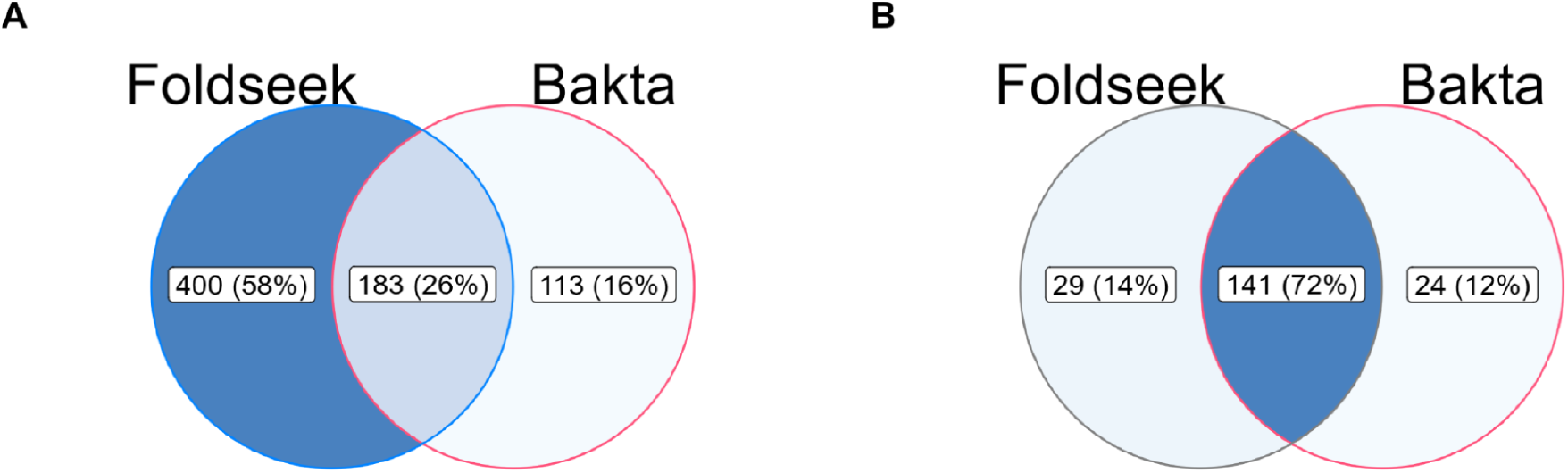
PFAM annotations were compared between Foldseek and Bakta. (A) All PFAMs annotations of predicted bacteriophage were pooled from all samples and compared for matches. (B) Pairwise comparisons were performed between Foldseek and Bakta PFAM annotations for each predicted CDS, to see if results from both methods agree. Matches are counted if at least one PFAM obtained from one method agrees with the other for a given CDS that was predicted by Bakta.

Finally, we clustered all proteins of the bacteriophage head, tail, capsid or baseplate categories that the Foldseek annotations identified from all putative bacteriophage by structural alignments (Figure 8). Near perfect TM scores within most clusters show that the same putative best structural homolog was often seen in samples widely separated by time, suggesting that similar bacteriophage genomic sequences were being captured on different collection dates. Unfortunately however, relating bacteriophage in our samples to known families of bacteriophage based on the cluster representative was not possible due to the lack of bacteriophage specific taxonomic information in the UniProt entries and the high sequence divergence precluded phylogenetic inference. However, future investigations considering factors such as the morphology of bacteriophage structures, bacteriophage hosts and functional proteins to classify our bacteriophage can be done with the annotation methods we highlight in this study.

**FIG 8.**
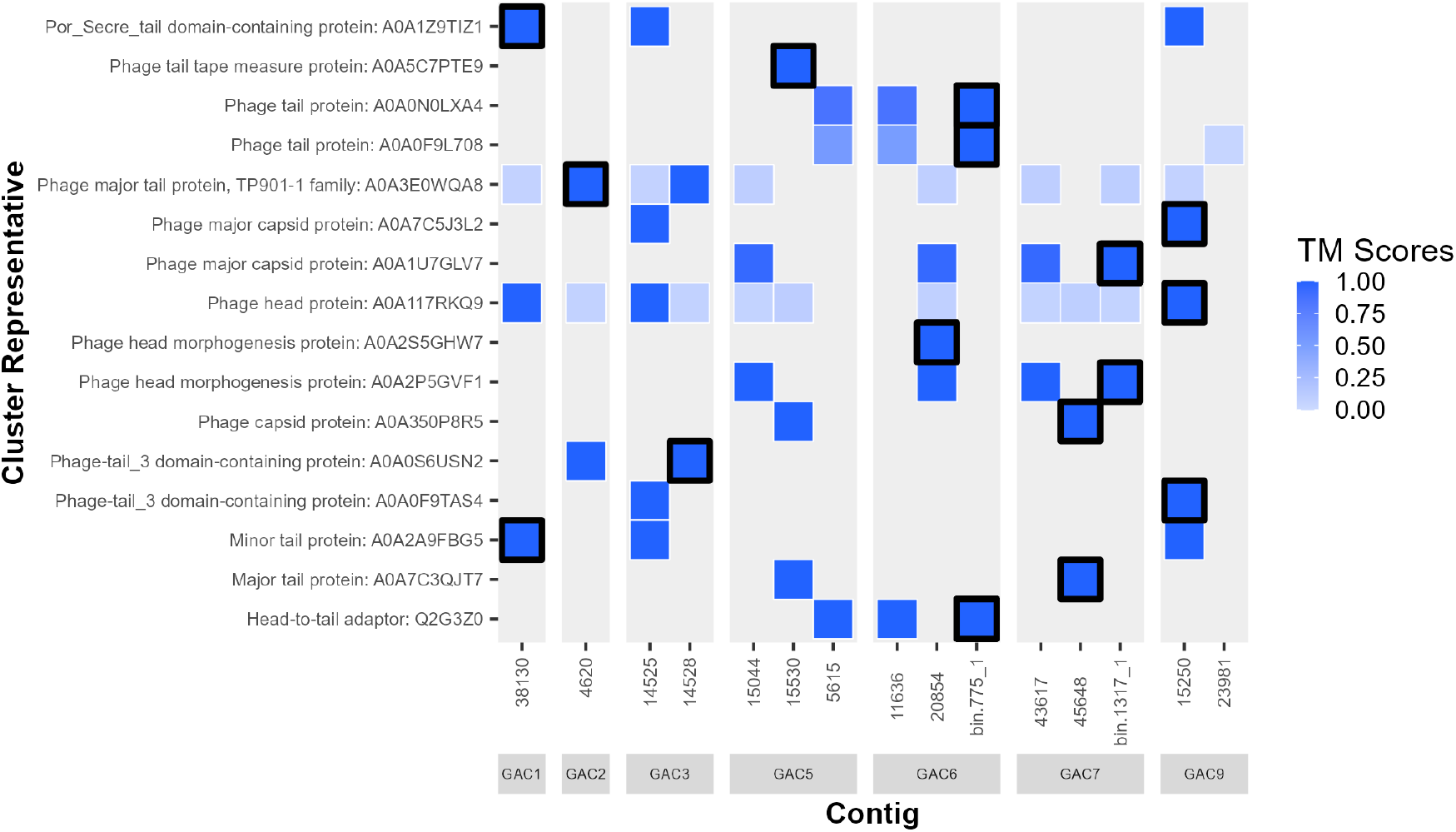
Structural bacteriophage proteins, which are often used for classification of bacteriophage, were identified via Foldseek and were shared between putative bacteriophage across samples. All predicted proteins structures belonging to recurring putative bacteriophage ACCs were clustered using Foldseek, and each cluster containing representatives that were annotated by Foldseek to be head, capsid, baseplate or tail related proteins had its representative (outlined black) structurally aligned to its members to generate a TM-score. Most clusters have members that strongly resemble its representative, having near perfect alignment scores close to 1 for the entire cluster. Clusters are named according to the UniProt recommended name for the representative protein structure, followed by its accession number

## CONCLUSIONS

We developed an integrated approach to characterize the gene content of unique microbial communities. We start by producing ultra-high molecular weight DNA from 10 GAC samples, which was subsequently sequenced on OXford Nanopore MinION platform. Using our custom workflow, we generated 5432 high quality and circular metagenomic assemblies in total with 227 being putative bacteriophage. However, annotation of these assemblies with Bakta, a state of the art sequence homology annotation method, left the vast majority of predicted genes as hypothetical for the ACCs, especially for putative bacteriophage. We demonstrate that using a combination of Colabfold, a protein structure prediction tool, and Foldseek, a homology search tool, can greatly enrich and improve annotations of assemblies that have high sequence divergence to the anything in current nucleotide or protein databases. With this approach, more comprehensive characterizations of microbial communities living in oil refinery wastewater can be generated, which would significantly aid in our current understanding the processes involved in NA biodegradation with further analyses.

## MATERIALS AND METHODS

### High Molecular Weight DNA Extraction and Library Preparation

One key component of producing circular assemblies was the preparation of ultra high molecular weight DNA for sequencing. Prior to extraction, GAC samples were collected and frozen at -80°C. Approximately 10 g of GAC and associated biofilm was added a 50 mL falcon tube, followed by 10 mL of lysis buffer (10 mM Tris-HCl, 100 mM NaCl, 25 mM EDTA, 0.5% (w/v) SDS) and 100 *μ*L of lysozyme (25 mg/mL). The sample and buffers were mixed by slowly rotating the tube while trying to minimize granule movement, and then incubated for 1 hour at 37°C. After 1 hour, 5 *μ*L of RNase A (20 mg/mL) was added, the sample was mixed gently, then incubated for 1 hour and 30 minutes at 57°C. Finally, 100 *μ*L of Proteinase K (800 units/mL) was added to the tube, gently mixed, and the sample was incubated for another 1 hour and 30 minutes at 57 °C.

The lysate was decanted into a new 50 mL falcon tube, without transferring the carbon granules. Then, 1 volume of 25:24:1 phenol:chloroform:isomayl alcohol was added to the lysate, the mixture was rocked gently for 8 minutes, then spun at 3000 x rcf for 3 minutes. The aqueous phase was transferred to a new 50 mL falcon tube using wide bore pipette tips. This process was repeated twice with 1 volume of chloroform instead of phenol:chloroform:isomayl alcohol, for a total of 2 chloroform washes.

To precipitate the DNA, 1/10 volume of 3 M sodium acetate at pH 4.5 was added, followed by 2 volumes of ice cold 100% ethanol. DNA that precipitated immediately was spooled out into an eppendorf tube with a sterile pasteur pipette that had been melted into a hook. The DNA was washed twice with nuclease free 75% ethanol and resuspended in 500 *μ*L to 1 mL of Tris buffer (10 mM, pH 8) overnight, depending on the size of the pellet.

Prior to library preparation, 60 *μ*L of the DNA sample was added to 60 *μ*L of 3% PVP 360000 solution (1.2 M NaCl, 20 mM Tris-HCl, pH 8) and mixed thoroughly by inverting. The mixture was spun at 10000 x rcf for 30 minutes at RT to preferentially precipitate high molecular weight DNA. The supernatant was discarded, then the DNA pellet was washed twice with 75% nuclease free ethanol. The DNA was resuspended in 60 *μ*L of Tris buffer (10 mM, pH 8).

The DNA library was prepared using the ONT LSK109 or LSK110 kit according to the manufacturers protocol, with the following changes. The repair and end prep step was extended to 15 minutes at 20°C and 15 minutes at 60°C instead of the recommended time of 5 minutes at each temperature. Additionally, Omega Biotek Mag-Bind beads were used in place of Ampure XP beads. The libraries were sequenced on a MinION, with 9.4.1 flow cells for libraries prepared with the LSK109 kit or LSK110 kit.

### Assembly

Raw reads were basecalled with Guppy v6.3.8 (17) using the arguments ‘-c dna_r9.4.1_450bps_sup.cfg –min_qscore 7’. Basecalled reads were checked for length and q-score using PycoQC (v2.5.2) (18) to determine a cutoff for lower quality data to discard. The minimum length was always greater than 500 nt with a minimum read q-score of 7. Once a cutoff was determined, NanoFilt (v2.8.0) (19) was used to filter the reads with the length and q-score parameters set to the cut off, as well as the argument “–headcrop 50”. The filtered reads were then assembled with Flye (v2.9-b1768) (20) in ‘–nano-hq –meta’ mode. After the initial assembly, additional assemblies were yielded using a secondary assembly pipeline. Briefly, reads for a given sample were aligned to uncircularized contigs obtained from the same sample with Minimap2 v2.24 (21) and were binned using MetaBAT2 v2.12.1 (22). Reads aligned to a bin were filtered using Gerenuq v0.2.3 (10) on default settings to keep only alignments over 1000 bp, with a score of 1 and at least 50% identity. For each bin, Gerenuq filtered reads were passed on to Flye with the genome size set to the total size of the bin. Only assemblies that had an estimated coverage of at least 10 and that were tagged as circularized by Flye were extracted from the assembly graphs as a GFA file and passed on to the polishing pipeline.

### Polishing

Before polishing, all metagenomic reads from a given sample were aligned to each assembly obtained from the same sample using Minimap2. Mapped reads were filtered with Gerenuq to keep alignments that are at least 1000 bases long, with a score of at least 1 and at least 90% identity. Then, draft GFA assemblies were polished with Minipolish v0.1.2 (23) using the Gerenuq filtered reads that were converted to fasta format. In order for Minipolish to accept Flye assemblies however, minor changes were made to the file format of Flye’s outputs namely by changing converting the GFA file format to GFA2 with GFAKluge (24) and by adding “l” as a suffix to sequence names. Additionally, Minipolish was run with the ‘–skip-intial’ argument as the initial step requires non-standard data unique to the Miniasm assembler. The polished assemblies were converted to fasta format, then underwent a second round of polishing with Medaka v1.6.1 (25) using the same Gerenuq filtered reads. To assess the coverage of polished assemblies, gerenuq filtered reads were mapped back to the polished assemblies and put through Mosdepth v0.3.3 (26) to calculate coverage by 1000 base windows. Polished assemblies that were greater than 1 mb in size were checked for completeness and contamination using Checkm v1.2.2 (27), and those that had greater than 90% completeness and less than 10% contamination were used as benchmarks for assessing the general quality of predicted bacteriophage assemblies.

### Annotation and bacteriophage Prediction

Bakta v1.5.1 (6) with the ‘–complete’ argument was used to annotate the polished assemblies. The assemblies were subset further based on Bakta annotations assemblies that contained ribosomal related proteins were discarded, and those that also had annotated bacteriophage related proteins were passed to INHERIT (11) to identify potential bacteriophage genomes using their pre-trained model. From each bacteriophage predicted by INHERIT, all amino acid sequences from Bakta annotations were passed to Colabfold v1.3.0 (8) with the arguments ‘–amber –templates –num-recycle 3 –use-gpu-relax’ to predict the structure of each protein. For each protein that had a structure prediction, the rank 1 model was taken and queried against the Alphafold/Uniprot database using Foldseek v90b (9) with the ‘easy-search’ function. Functional annotations including KEGG and GO terms were retrieved using Uniprot’s API by querying only the best Foldseek hits, which are filtered for an e-value greater than 1e-10, for each predicted protein structure. Pathway information for KEGG KO’s retrieved with Foldseek was obtained using the KEGGREST 1.38.0 package (28).

### Singleton and Recurring Assemblies

Singleton and recurring circularized and polished assemblies were determined by performing an all versus all BLAST. A pair of assemblies from different samples were counted as identical if they had a percent identity of 98%, a query coverage that is within 10% of the query length, and a query length that is at least 90% of subject length. Each set of ACCs that were considered singleton or recurring were subset further if they were predicted bacteriophage. To observe the presence of structural bacteriophage proteins across samples, the total set of proteins from each recurring predicted bacteriophage ACC was collected and clustered with Foldseek using the greedy set cover algorithm, and alignments where 80% of the sequence is covered by the alignment are kept. The clusters were subset based on keywords in their representative’s Foldseek annotations “head/tail/capsid/plate”. For each cluster in which the representative is a putative head, tail, capsid or baseplate related protein, the protein structure of the cluster representative was aligned with its members to obtain a TMScore using Foldseek.

### Data and Code Availability

Raw read data was deposited to the European Nucleotide Archive, under study accession PRJEB49151. Code used in the methods and figures presented in this manuscript is available at our Github repository.

## SUPPLEMENTAL MATERIAL

TABLE S1. INHERIT prediction scores for each ACC that contained bacteriophage proteins in the initial annotation with Bakta.

FIGURE S1. The coverages across the length of putative bacteriophage ACCs in 1% bin windows, ordered from highest standard deviation to lowest. Bacteriophage ACCs denoted with ** contain full length alignments with 90% sequence identity to the reference. Labels in red indicate those ACCs with SD/cov ratios greater than 1.

## ACKNOWLEDGMENTS

We thank Dave Edgell and his lab for sharing their lab space. We also would like to thank Mitacs for funding the project with the Accelerate Fellowship grant. Finally, we would like to thank Dr. Martin Flatley for his support with the collection of samples for this study.

